# Local and non-local impacts of intra-hexamer interactions on temperature compensation of KaiC

**DOI:** 10.64898/2026.06.12.727564

**Authors:** Kanta Kondo, Yoshihiko Furuike, Kota Horiuchi, Yasuhiro Onoue, Eiki Yamashita, Shuji Akiyama

**Affiliations:** Research Center of Integrative Molecular Systems (CIMoS), Institute for Molecular Science, National Institute of Natural Sciences, 38 Nishigo-Naka, Myodaiji, Okazaki 444-8585, Japan; Molecular Science Program, Graduate Institute for Advanced Studies, SOKENDAI, 38 Nishigo-Naka, Myodaiji, Okazaki 444-8585, Japan; Institute for Protein Research, The University of Osaka, Suita, 565-0871, Japan

## Abstract

A hexameric clock protein KaiC exhibits a 24-hour phosphorylation cycle with a unique property termed temperature compensation. The period is kept constant over physiological temperatures through compensatory coordination of underlying elementary reactions. The temperature-compensated ATPase activity of KaiC is one such key reactions that potentially contribute to maintaining a constant circadian period. We identified four amino acid residues responsible for the temperature compensation in an N-terminal ATPase domain of KaiC. D82 and K172 were located in a primary site, and the ATPase activity of each alanine mutant showed a positive correlation with rising temperature. N62 and E69 constituted a secondary site, where each alanine replacement resulted in a negative correlation with the temperature. The primary site exerts a compensatory regulation over the ATPase cycle locally within the N-terminal domain. The secondary site prevents the ATPase activity from becoming over-compensated by suppressing another compensatory regulation mediated through a non-local interaction with a C-terminal domain of KaiC. Therefore, any imbalance between the local and non-local compensatory regulations in KaiC affects the temperature dependence of its phosphorylation rhythm.

## Introduction

Circadian clocks are endogenous timing systems that allow organisms to adapt to daily environmental changes caused by Earth’s rotation. The circadian clocks have been identified in diverse organisms, including bacteria (Ishiura *et al*, 1998), fungi (McClung *et al*, 1989), plants (Strayer *et al*, 2000), insects (Reddy *et al*, 1984), and mammals (King *et al*, 1997), and they share three essential properties. First, the circadian clock generates an autonomous oscillation with a period of approximately 24 h even under constant conditions (self-sustainment). Second, the phase of the clock system can be shifted and entrained by external stimuli such as light and temperature (synchronization). Third, the period length of the circadian rhythms is kept almost constant over a range of physiological temperatures (temperature compensation). These three properties endow the circadian clocks with robustness and adaptability to the environmental fluctuations, enabling organisms on Earth can suitably control their physiological processes throughout day/night cycles.

Temperature compensation is the property specific to the circadian clocks, indicating a unique relationship between temperature and reaction rates. The temperature dependence of biochemical reactions is often evaluated using the temperature coefficient (*Q*_10_), a factor by which the reaction is accelerated by increasing the ambient temperature by 10°C. The *Q*_10_ values of the frequency (1/period) of the circadian clocks are mostly in the range of 0.9–1.1 (Hu *et al*, 2021; Joshi *et al*, 2022; Maeda *et al*, 2024; Nakajima *et al*, 2005; Shinohara *et al*, 2017), while general reactions including Belousov-Zhabotinsky oscillator often exhibit *Q*_10_ of 2–3 (Elias *et al*, 2014; Ruoff, 1995). This means that the period length of the clock system is almost temperature-independent, although it should consist of temperature-dependent elementary steps. The absolute slowness suggests that the clock system possesses a much higher activation energy barrier (*E*_A_) and is therefore highly temperature-dependent. To solve this apparent inconsistency between the slowness and temperature compensation, it is necessary to investigate the effect of the temperature on the kinetics and structure of clock components.

KaiC is a core clock protein responsible for the temperature compensation of the circadian clock system in cyanobacterium *Synechococcus elongatus* PCC 7942 (Furuike *et al*, 2022c; Nakajima *et al*., 2005; Terauchi *et al*, 2007). KaiC consists of an N-terminal (CI) adenosine triphosphatase (ATPase) domain and a C-terminal (CII) kinase/phosphatase domain (Fig. 1A), and forms a double-ring hexamer upon binding one molecule of adenosine triphosphate (ATP) in every CI–CI interface and CII–CII interface (Pattanayek *et al*, 2004). When KaiC is mixed with KaiA and KaiB in a test tube (KaiABC system), KaiC exhibits a temperature-compensated cycle between its phosphorylated and dephosphorylated states (P-cycle) (Nakajima *et al*., 2005; Nishiwaki *et al*, 2007; Rust *et al*, 2007) via the interaction of the Kai proteins (Akiyama *et al*, 2008; Kageyama *et al*, 2006; Snijder *et al*, 2017; Tseng *et al*, 2017) and the hydrolysis of ATP by KaiC (Terauchi *et al*., 2007). Previous studies showed that the frequency of the P-cycle is proportional to the ATPase activity of KaiC (Abe *et al*, 2015; Terauchi *et al*., 2007). The ATPase activity of KaiC is robustly temperature-compensated (*Q*_10_ ∼ 1) (Furuike *et al*., 2022c; Terauchi *et al*., 2007), even though it is approximately 10^3^ to 10^4^-fold lower than other ATPases such as kinesin (Cochran *et al*, 2006), myosin (Giese & Spudich, 1997), and F_1_-ATPase (Ackerman *et al*, 1987). These findings suggest that the essential properties of the cyanobacterial clock system—slowness and temperature compensation—are closely related to the ATPase activity of KaiC (Abe *et al*., 2015; Furuike *et al*., 2022c).

**Figure 1.**
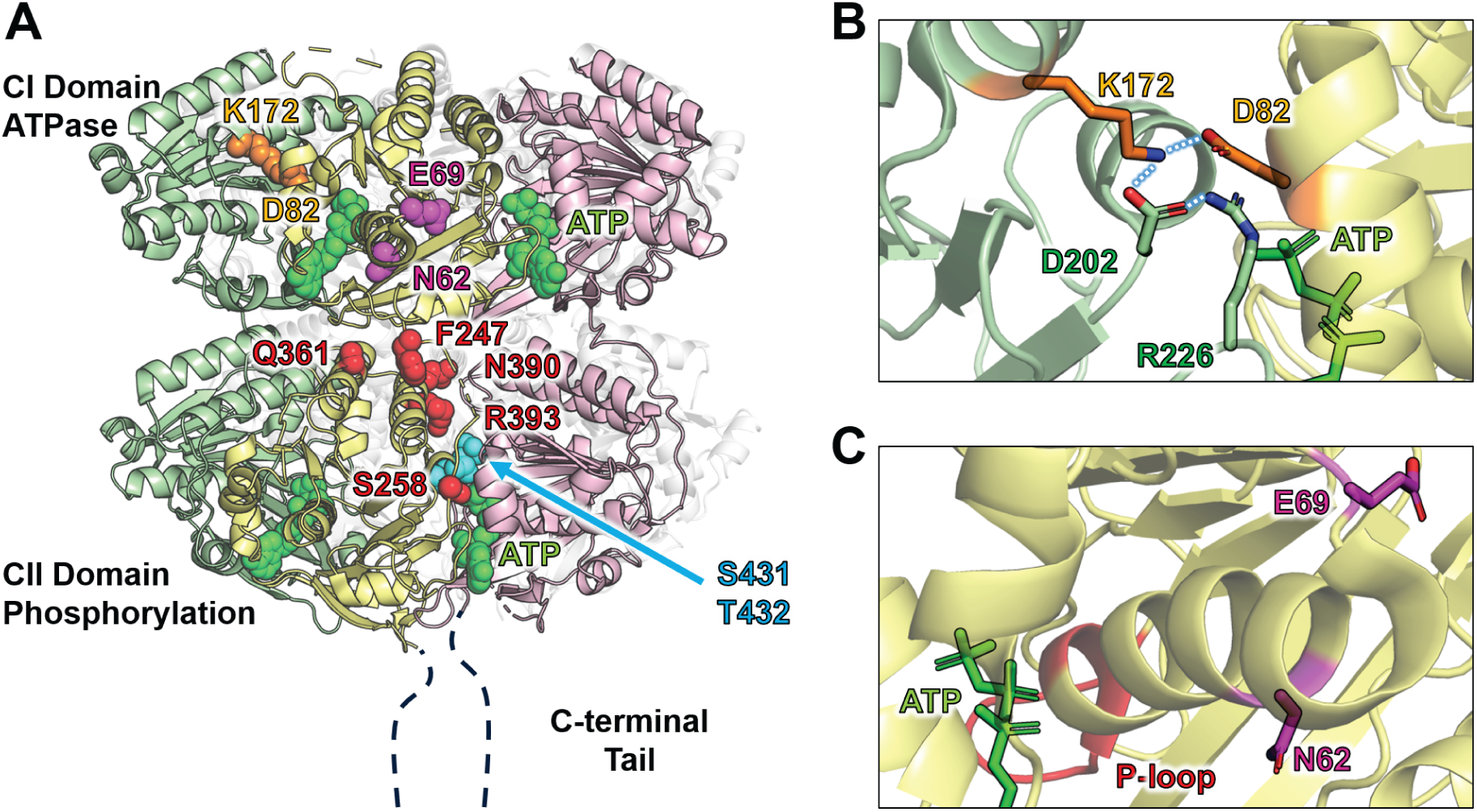
KaiC as a key component in the temperature compensation of cyanobacterial circadian rhythms. (**A**) Amino acid residues involved in the temperature compensation mapped onto the crystal structure of KaiC (PDB ID: 7DXQ) (Furuike *et al*, 2022b). KaiC forms a double-ring hexamer, in which each of CI and CII domains form a ring structure by incorporating one ATP molecule (green spheres) at every protomer-protomer interface. Two phosphorylation sites, S431 and T432 (cyan spheres), are located in the CII–CII interface. Four amino acid residues in the CI domain identified through this study are highlighted by orange (D82 and K172) or magenta sphere models (N62 and E69). All other amino acid residues (F247, S258, Q361, N390, and R393) identified previously (Furuike *et al*., 2022c; Murayama *et al*., 2011) are located in the CII domain and highlighted as red sphere models. (**B**) A zoomed-in-view of D82 and K172 in an interface between two CI protomers (yellow and green) (PDB ID: 7V3X). Dotted blue lines represent hydrogen bonds. (**C**) A zoomed-in-view of N62 and E69 (PDB ID: 7DXQ). The two residues are located in a C-terminus region of the α_2_-helix.

Several efforts have been made to isolate temperature-sensitive mutants of KaiC. An earlier study (Murayama *et al*, 2011) showed that a KaiC mutant with the R393C substitution (KaiC^R393C^) shows a reasonably temperature-compensated ATPase activity, but exhibits a temperature-sensitive P-cycle in the presence of KaiA and KaiB. A subsequent study (Furuike *et al*., 2022c) reported a series of CII variants of KaiC, in which both the ATPase activity and P-cycle are accelerated in a temperature-dependent manner. All the mutation sites identified previously as impairing the temperature compensation were confined to the CII domain (Fig. 1A). Therefore, it remains unclear what role the CI-ATPase domain plays in terms of the temperature compensation and how it relates to the CII domain.

In this study, we report four amino acid residues in the CI domain (D82, K172, N62, and E69 in Fig. 1A), where each alanine substitution impairs the temperature-compensatory nature of the ATPase activity. The temperature-sensitive CI mutants were largely categorized into two groups based on their characteristics and locations. One is located at the CI–CI interface (Fig. 1B) and shows increased activities with rising temperature (*Q*_10_ > 1, positive temperature-dependence), while the other is situated around the C-terminal region of the α_2_-helix (Fig. 1C) and exhibits reduced activities at high temperatures (*Q*_10_ < 1, negative temperature-dependence). The CI–CI interface exerts a local compensatory control over the active site, whereas a region near the C-terminus of the α_2_-helix suppresses a non-local compensatory regulation mediated by the CII domain. The ATPase activity of an N-terminal fragment consisting of the CI domain (ΔKaiC) is mildly temperature-compensated on its own (*Q*_10_ ∼ 1.5), but this inter-domain regulation from the CII to CI domains is necessary for the perfect temperature compensation observed in full-length KaiC (*Q*_10_ ∼ 1). The temperature-compensated ATPase activity of KaiC is one of the key factors underlying the temperature compensation of the P-cycle, and it confers the perfect temperature compensation to the P-cycle through complementary interactions with other Kai proteins.

## Results and Discussion

### ATPase activity of temperature-sensitive KaiC mutants

At the standard temperature of 30°C, the ATPase activity of wild-type KaiC (KaiC^WT^) was determined to be 12.9 ± 0.8 d^−1^ and nearly constant over the range of 25°C to 35°C (Fig. 2A). An apparent activation energy of KaiC^WT^ (*E*_app_^WT^) was estimated by Arrhenius-plot analysis to be 0.38 kcal mol^−1^ (Fig. 2B). The *Q*_10_ value of the ATPase activity (*Q*_10atp_^WT^) at 30°C was calculated to be 1.02 ± 0.15 by substituting *E*_app_^WT^ into Eq. 1 (Furuike *et al*, 2016) (details in Methods), which means that the ATPase activity of KaiC^WT^ is almost temperature-independent as previously reported (Furuike *et al*., 2022c; Terauchi *et al*., 2007).

**Figure 2.**
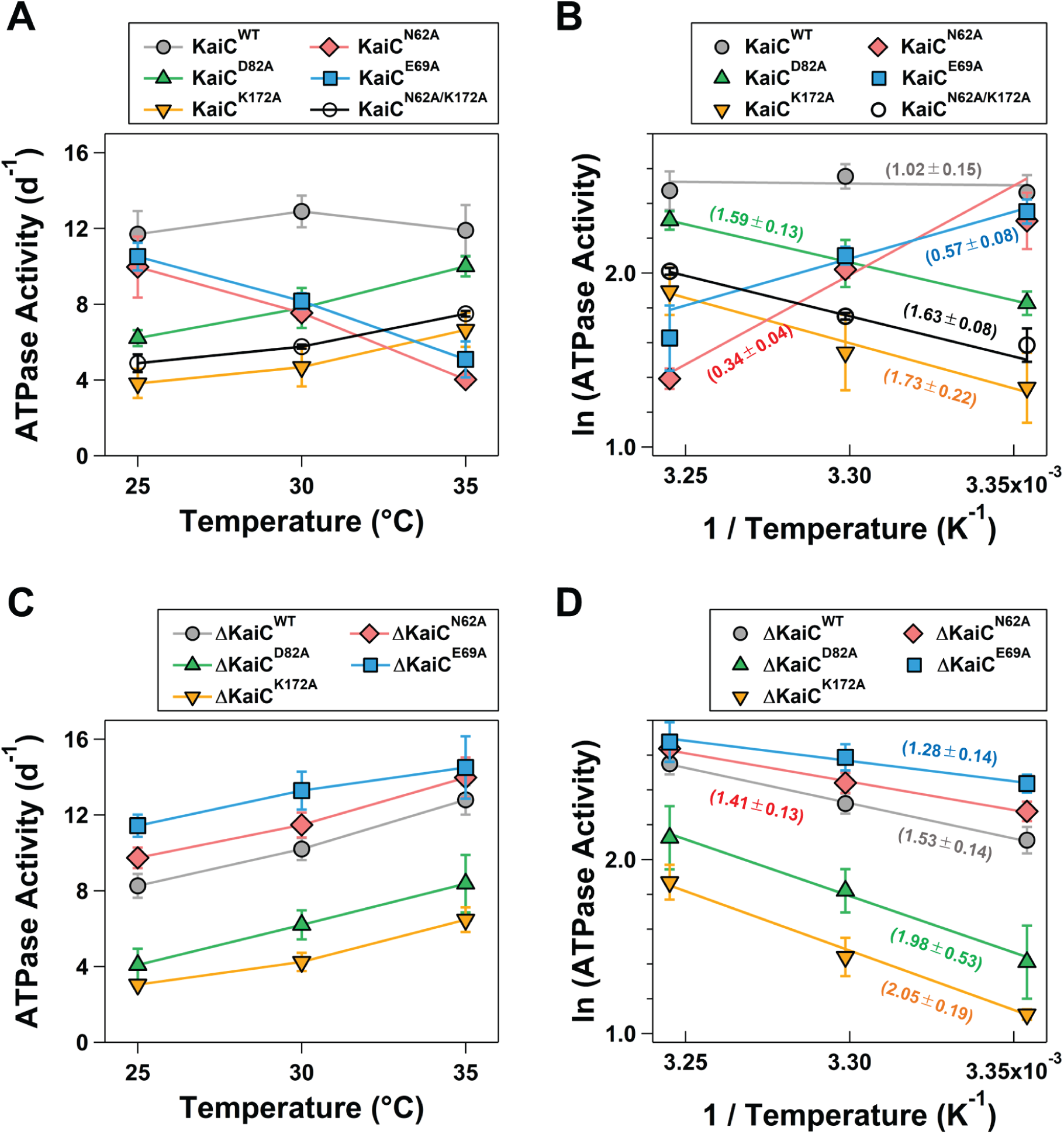
Temperature dependence of ATPase activities measured at 25°C, 30°C, and 35°C. (**A**) ATPase activities of full-length KaiCs. (**B**) Arrhenius-plot analysis of full-length KaiCs. Values in parentheses represent *Q*_10atp_. (**C**) ATPase activities of N-terminal fragments consisting of the CI domain (ΔKaiCs). (**D**) Arrhenius-plot analysis of ΔKaiCs. Plots and error bars indicate mean ± standard deviation (s.d.) from three replicates measured at different days, except for ΔKaiC^WT^ (five replicates).

A KaiC mutant with the D82A substitution (KaiC^D82A^) and KaiC^K172A^ exhibited positively temperature-dependent ATPase activities. At 25°C, the ATPase activity of KaiC^D82A^ was nearly half that of KaiC^WT^ (6.2 ± 0.4 d^−1^), but at 35°C, it reached an activity indistinguishable from KaiC^WT^ (10.0 ± 0.5 d^−1^) (Fig. 2A). The *Q*_10atp_^D82A^ value was estimated to be 1.59 ± 0.13 (Fig. 2B), which is comparable to that previously reported for the CII variant (*Q*_10atp_^Q361E^ = 1.7) (Furuike *et al*., 2022c). While KaiC^K172A^ showed reduced activities compared to KaiC^D82A^ at each temperature (Fig. 2A), it exhibited the same temperature dependence (*Q*_10atp_^K172A^ = 1.73 ± 0.22) as KaiC^D82A^ (Fig. 2B).

In contrast, the ATPase activities of KaiC^N62A^ and KaiC^E69A^ followed negative temperature-dependences. At 25°C, KaiC^N62A^ showed an activity equivalent to that of KaiC^WT^ (10.0 ± 1.6 d^−1^), whereas at 35°C, its ATPase activity was approximately 60% lower than that of KaiC^WT^ (4.0 ± 0.2 d^−1^) (Fig. 2A). The *Q*_10atp_^N62A^ value was estimated to be 0.34 ± 0.04 (Fig. 2B). The ATPase activity of KaiC^E69A^ was also decreased with rising temperature and *Q* ^E69A^ was determined to be 0.57 ± 0.08 (Fig. 2A and 2B).

### Functional reversibility of KaiC^N62A^ and KaiC^E69A^ after high-temperature treatment

Enzymes often decrease their activities due to thermal denaturation at temperatures higher than their optimal range. Therefore, two possibilities can be considered regarding the negative temperature-dependence observed in KaiC^N62A^ and KaiC^E69A^. First, the reduction of *Q*_10atp_ to approximately 0.4 may be due to unfolding of approximately 60% of the CI variants at the high temperature of 35°C. Second, both KaiC^N62A^ and KaiC^E69A^ are over-compensated mutants.

To examine these two possibilities, we investigated the reversibility of the ATPase activity under a temperature cycle. As shown in Fig. 3A, the ATPase assay of KaiC^WT^ was initiated at 25°C at an incubation time (IT) of zero. At IT = 9 h, the incubation temperature was elevated from 25 to 35°C and kept at 35°C for 9 h. At IT = 18 h, the temperature was lowered from 35 to 25°C and kept at 25°C for 9 h. Despite being under the temperature cycle, KaiC^WT^ produced ADP monotonously over time (Fig. 3A). KaiC^N62A^ and KaiC^E69A^ were subjected to the same temperature cycle as KaiC^WT^. In contrast to KaiC^WT^, the ADP production by KaiC^N62A^ was dramatically decreased at 35°C (2.4 ± 0.1 d^−1^, IT = 9–18 h) compared to at 25℃ (7.9 ± 0.3 d^−1^, IT = 0–9 h) (Fig. 3B). The degree of reduction is nearly comparable to *Q*_10atp_^N62A^ (0.34 ± 0.04). When the incubation temperature was lowered to 25°C (IT = 18–27 h), KaiC^N62A^ restored the ADP production to a level similar to that observed during the IT of 0–9 h. KaiC^E69A^ exhibited the similar tendency (Fig. 3B). The ADP production by KaiC^E69A^ was decreased at 35°C (4.2 ± 0.1 d^−1^, IT = 9–18 h) but increased after decreasing the temperature to 25°C (11.5 ± 0.7 d^−1^, IT = 18–27 h). The recovery of the ADP-production ability following the high-temperature treatment suggests that the decreases in *Q*_10atp_^N62A^ and *Q*_10atp_^E69A^ to approximately 0.4 is not due to unfolding of approximately 60% of KaiC. Thus, both KaiC^N62A^ and KaiC^E69A^ are over-compensated mutants: they possess excessively enhanced compensatory regulation of suppressing the activity increase caused by temperature elevation.

**Figure 3.**
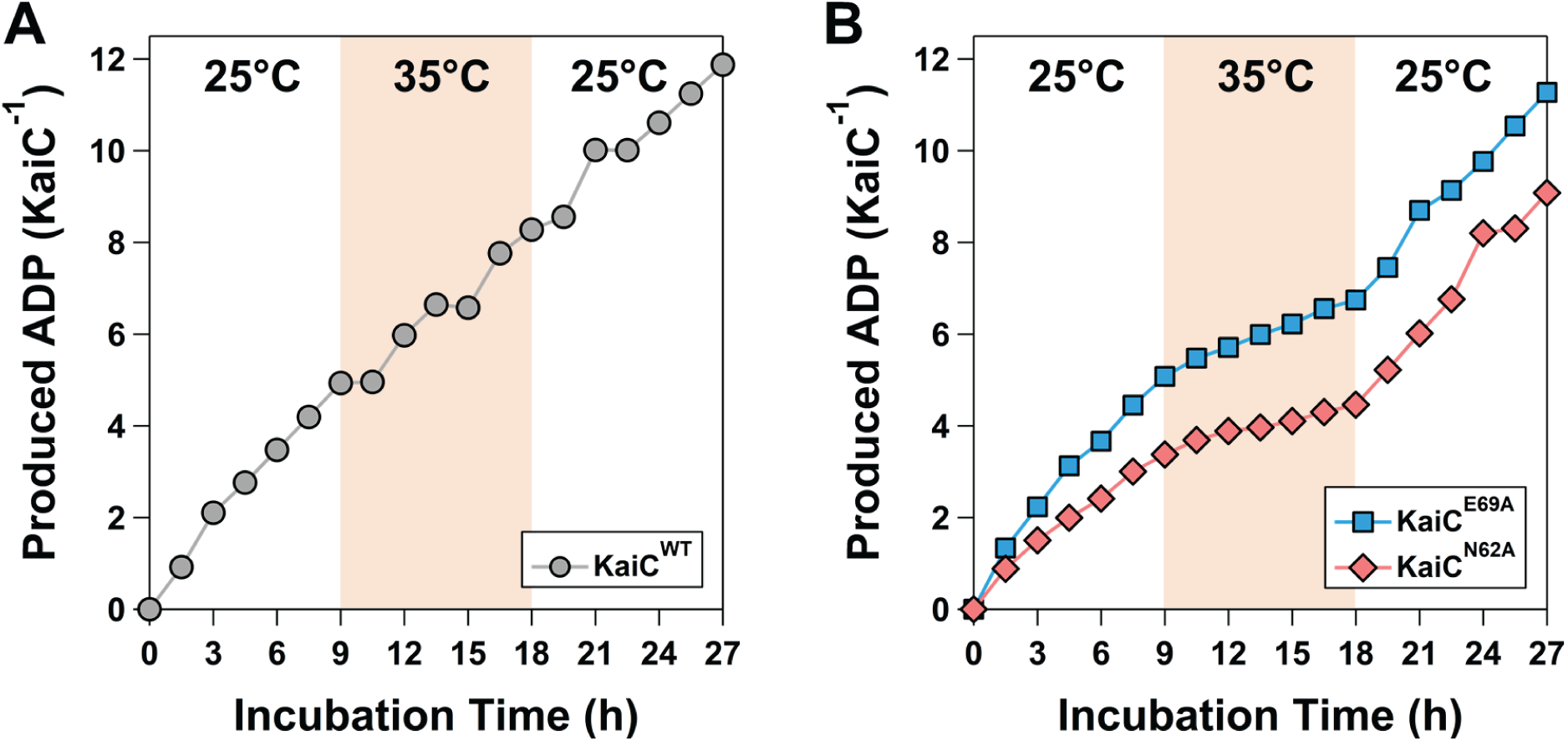
Reversibility of the ATPase activities. ADP productions by (**A**) KaiC^WT^, and by (**B**) KaiC^N62A^ and KaiC^E69A^ under temperature cycling from 25℃ to 35℃ and back to 25℃. Vertical axis indicates the number of ADP produced by one KaiC molecule in a monomer unit. Horizontal axis indicates the incubation time after the temperature cycle started.

### Local and non-local compensatory regulations in achieving robust temperature compensation in KaiC

As shown in Fig. 1A, D82 and K172 are located in close proximity to each other at the protomer-protomer interface of the CI domain (site-I). Similarly, N62 and E69 are positioned at a distance of just 9 Å, but they are located in the outer-radius side of the hexameric ring (site-II). To investigate the effect of the CII domain on these two sites, we prepared fragments of the CI domain without (ΔKaiC^WT^) or with each of the four amino acid substitutions (ΔKaiC^D82A^, ΔKaiC^K172A^, ΔKaiC^N62A^, and ΔKaiC^E69A^).

The ATPase activity of ΔKaiC^WT^ was 8.3 ± 0.6 d^−1^ at 25°C (Fig. 2C), which accounted for approximately 70% of the ATPase activity of the full-length form (Fig. 2A) (Abe *et al*., 2015; Furuike *et al*, 2022a; Terauchi *et al*., 2007). The activity was increased with rising temperature, and resultant *Q*_10atp_^ΔWT^ was 1.53 ± 0.14 (Fig. 2D). Although ΔKaiC^WT^ is not perfectly temperature-compensated like full-length KaiC^WT^, its temperature dependence yet remains very low compared to general enzymes with *Q*_10_ as high as a 2–3 (Elias *et al*., 2014). This result clearly demonstrates that the CI-ATPase is locally and moderately temperature-compensated even when isolated (*Q*_10atp_^ΔWT^ = 1.53 ± 0.14), and requires some non-local communications with the CII domain to achieve the perfect temperature compensation (*Q*_10atp_^WT^ = 1.02 ± 0.15).

We found that the site-I mutations are effective even in the absence of the CII domain. The ATPase activity of ΔKaiC^D82A^ increased more sharply with rising temperature than that of ΔKaiC^WT^, and consequently *Q*_10atp_^ΔD82A^ was estimated to be 1.98 ± 0.53 (Fig. 2D). ΔKaiC^K172A^ also exhibited an enhanced positive temperature-dependence (*Q*_10atp_^ΔK172A^ = 2.05 ± 0.19, Fig. 2D). In terms of the temperature compensation, therefore, the site-I serves as a site that locally downregulates the positive temperature-dependence of the CI-ATPase, because *Q*_10atp_^ΔD82A^ (1.98 ± 0.53) and *Q*_10atp_^ΔK172A^ (2.05 ± 0.19) are notably greater than *Q*_10atp_^ΔWT^ (1.53 ± 0.14) (Fig. 2D).

Surprisingly, in the site-II mutants, the truncation of the CII domain resulted in a reversal of the temperature dependence. In contrast to full-length KaiC^N62A^, whose ATPase activity followed the negative temperature-dependence (Fig. 2A), the activity of ΔKaiC^N62A^ increased positively with increasing temperature (Fig. 2C). A similar result was also observed in ΔKaiC^E69A^. The resultant *Q*_10atp_^ΔN62A^ and *Q*_10atp_^ΔE69A^ values were 1.41 ± 0.13 and 1.28 ± 0.14, respectively (Fig. 2D), and they were similar to *Q*_10atp_^ΔWT^ (1.53 ± 0.14). This similarity in *Q*_10atp_ is also supported by the fact that crystal structures of ΔKaiC^WT^, ΔKaiC^N62A^, and ΔKaiC^E69A^ are essentially identical (Appendix Fig. S1 and Table S1). Thus, the site-II is heavily involved in the non-local compensatory regulation by the CII domain, whereas it plays a quite limited role in locally downregulating the temperature dependence of the ATPase activity within the CI domain.

To investigate the relationship between the two sites, we prepared a double mutant with the N62A and K172A substitutions (KaiC^N62A/K172A^) and measured the temperature dependence of its ATPase activity. Interestingly, KaiC^N62A/K172A^ exhibited a positively temperature-dependent ATPase activity (*Q*_10atp_^N62A/K172A^ = 1.63 ± 0.08) similar to KaiC^K172A^, and the negative temperature-dependence observed in KaiC^N62A^ was completely disappeared (Fig. 2A and 2B).

There are two types of regulatory models that explain above observations. In one model (model-A in Fig. 4), the site-I is involved in the ATPase cycle via a local pathway that provides the temperature-compensatory regulation within the CI domain (red ball-tipped arrows in Fig. 4A). Furthermore, there is another temperature-compensatory pathway that acts non-locally through the assistance of the CII domain (blue ball-tipped arrows in Fig. 4A). The non-local compensatory pathway runs near the site-II and is under the inhibitory control by the site-II to prevent the ATPase activity from becoming over-compensation (black inhibitory arrows in Fig. 4A). The site-I mutants of full-length KaiC are considered to be in a state where the local compensatory regulation by the site-I is significantly weakened (Fig. 4B). On the other hand, the site-II variants are interpreted as being in an over-compensated state due to the attenuation or loss of the inhibitory ability (Fig. 4C). Furthermore, ΔKaiC^WT^ is regarded as a state lacking only the non-local compensatory regulation (Fig. 4E), which accounts for why *Q*_10atp_^ΔKaiC^ increased and unchanged upon the mutations of the site-I (Fig. 4F) and site-II (Fig. 4G), respectively. In the model-A, an explanation is needed as to why the positive temperature-dependence associated with the site-I mutations appears in the site-I/II double mutants. One plausible assumption is that the compensatory regulation suppressed by the site-II is effective only when the CI-ATPase cycle is under the local compensatory control via the site-I (Fig. 4D).

**Figure 4.**
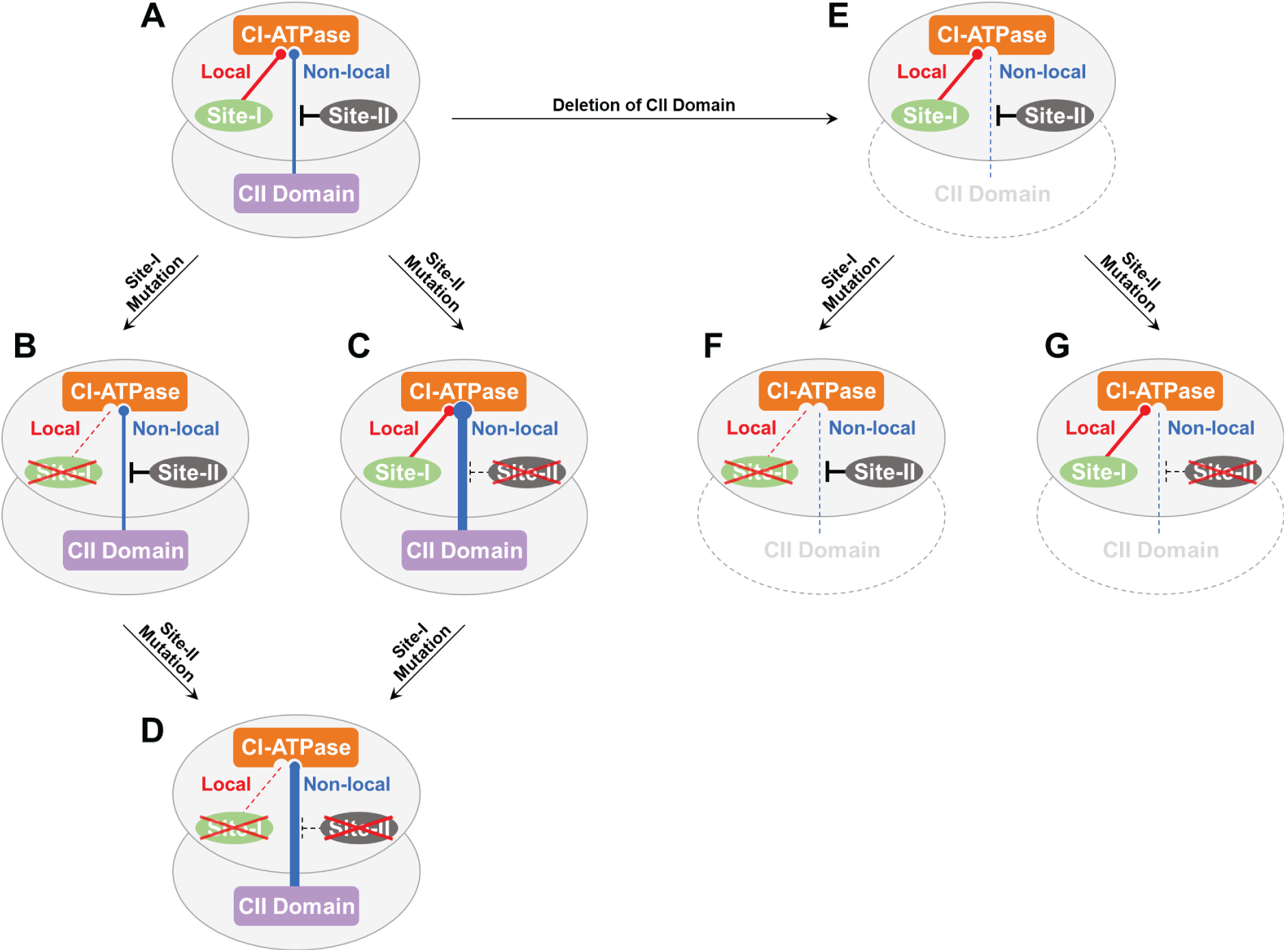
A conceptual illustration of local and non-local compensatory regulations in KaiC (model-A). A site-I acts on the CI-ATPase cycle through a local temperature-compensatory regulation (red-tipped arrows). A site-II inhibits (black inhibitory arrows) a non-local compensatory regulation from the CII to CI domains (blue-tipped arrows). (**A**) KaiC^WT^. (**B**) Site-I mutants of KaiC^WT^. The local compensatory regulation is absent or considerably weakened. (**C**) Site-II mutants of KaiC^WT^. The non-local compensatory regulation is excessively enhanced (thick blue-tipped arrow) by the release of inhibition by the site-II, resulting in the over-compensated CI-ATPase. (**D**) Site-I/II double mutant of KaiC^WT^. The non-local compensatory regulation is assumed to be effective only when the CI-ATPase cycle is under local compensatory control mediated by the site-I. (**E**) ΔKaiC^WT^. The deletion of the CII domain results in a loss or significant attenuation of the non-local compensatory regulation. (**F**) Site-I mutants of ΔKaiC^WT^. (**G**) Site-II mutants of ΔKaiC^WT^. The CI-ATPase activity is yet under some local compensatory regulation.

The other model differs from the model-A in that its non-local compensatory regulation acts via the site-I (model-B in Appendix Fig. S2). The site-I is a receiver of the non-local compensatory regulation from the CII domain and forwards it to the active site of CI-ATPase (blue ball-tipped arrows in Appendix Fig. S2A). This forwarding step is under the inhibitory regulation by the site-II to prevent the excessive compensation of the ATPase activity (black inhibitory arrows in Appendix Fig. S2A). Each of the site-I and site-II mutations can be explained in the same way as described above, regardless of whether they are under the full-length (Appendix Fig. S2B and S2C) or ΔKaiC background (Appendix Fig. S2E, S2F, and S2G). The model-B also explains why the site-I/II double mutants exhibited the same positive temperature-dependence as the site-I mutants (Appendix Fig. S2D).

### Phosphorylation cycle of temperature-sensitive KaiC mutants

To examine the effect of *Q*_10atp_ on the KaiABC system, we measured the P-cycles of KaiC^WT^ and its mutants in the presence of KaiA and KaiB. The P-cycle length of KaiC^WT^ was 24.4 ± 0.2 h (Fig. 5A) at 30°C, which corresponded to a P-cycle frequency (*f*_pc_) of 0.98 ± 0.01 d^−1^ (= 24 / period). Even when the temperature was raised by 5°C or 10°C from 30°C, the increase in *f*_pc_ was less than 10% of the result obtained at 30°C. Consequently, the *Q*_10_ value of the P-cycle frequency (*Q*_10fpc_^WT^) was estimated to be 1.08 ± 0.02 and was in good agreement with *Q*_10atp_^WT^ (1.02 ± 0.15).

**Figure 5.**
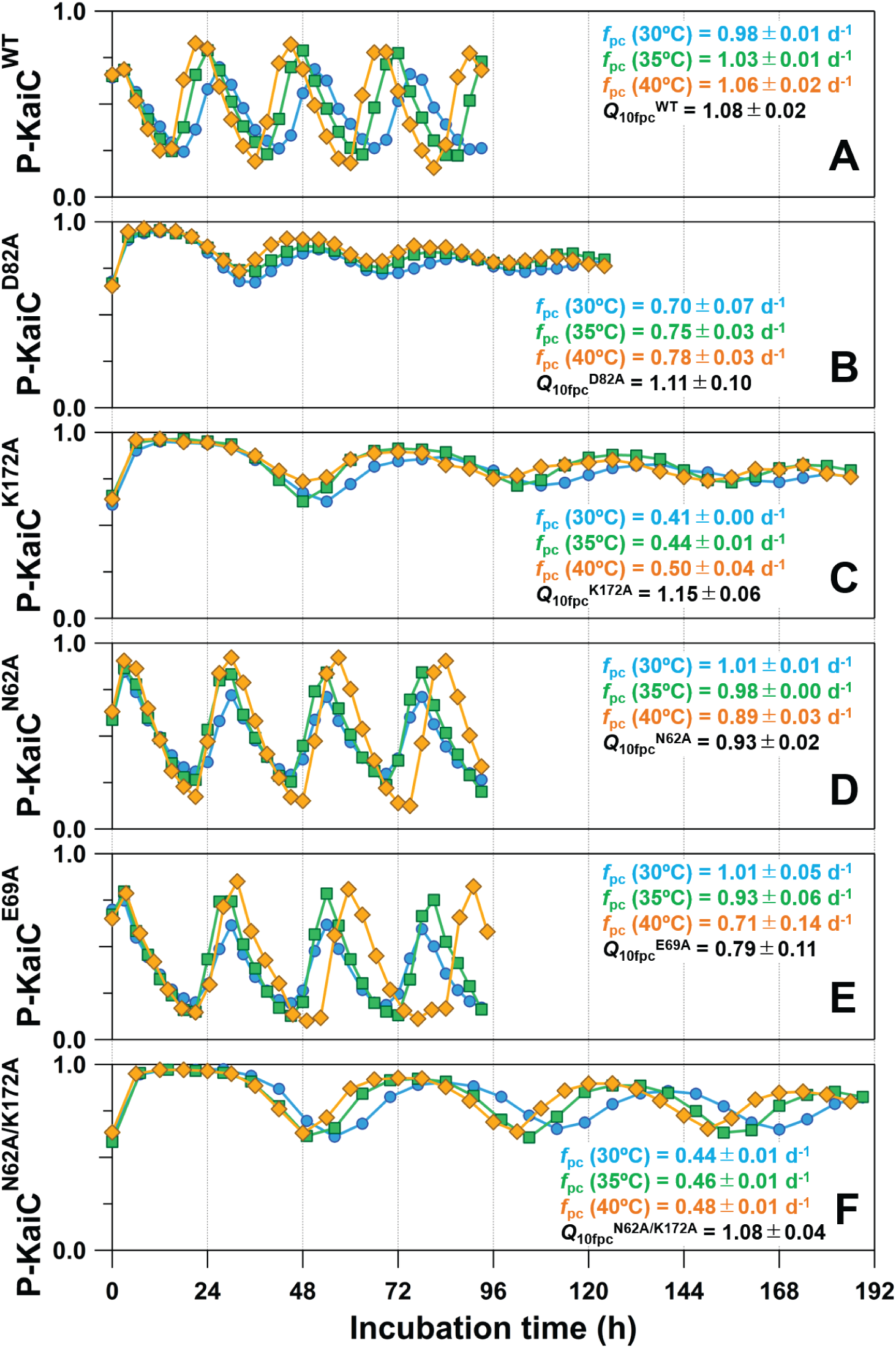
Rhythmic changes in the fraction of the phosphorylated KaiC (P-KaiC) at 30°C (cyan), 35°C (green), and 40°C (orange) in the presence of KaiA and KaiB. (**A**) KaiC^WT^, (**B**) KaiC^D82A^, (**C**) KaiC^K172A^, (**D**) KaiC^N62A^, (**E**) KaiC^E69A^, and (**F**) KaiC^N62A/K172A^. Each plot shows the representative result from three biological or technical replicates, with the exception of KaiC^E69A^ (six replicates). Frequencies of the phosphorylation cycle (*f*_pc_) are calculated by dividing 24 hours into the period length, and given as mean ± s.d. from three or six replicates. Temperature coefficients of *f*_pc_ at 30°C (*Q*_10fpc_) are determined by using Arrhenius-plot analysis and Eq. 1.

At the standard temperature of 30°C, the *f*_pc_ value of KaiC^D82A^ was 0.70 ± 0.07 d^−1^ (Fig. 5B) and lower than that of KaiC^WT^ (0.98 ± 0.01 d^−1^). This result is consistent with the observation that the ATPase activity of KaiC^D82A^ was lower than that of KaiC^WT^ at 30°C (Fig. 2A), because there is a positive correlation between *f*_pc_ and the ATPase activity (Abe *et al*., 2015; Furuike *et al*., 2022a; Terauchi *et al*., 2007). The *f*_pc_ value of KaiC^D82A^ was almost temperature-compensated (*Q*_10fpc_^D82A^ = 1.11 ± 0.10) in contrast to significant temperature-dependent acceleration of its ATPase activity (*Q*_10atp_^D82A^ = 1.59 ± 0.13). A similar tendency was observed also in KaiC^K172A^ (Fig. 5C); *f*_pc_ exhibited a less pronounced positive temperature-dependence (*Q*_10fpc_^K172A^ = 1.15 ± 0.06) compared to the ATPase activity (*Q*_10atp_^K172A^ = 1.73 ± 0.22).

The site-II mutants exhibited robust P-cycles with larger amplitude at every temperature. The *f*_pc_ value of KaiC^N62A^ at 30°C was 1.01 ± 0.01 d^−1^ (Fig. 5D) and nearly identical to that of KaiC^WT^ (Fig. 5A). In contrast to KaiC^WT^, however, *f*_pc_ of KaiC^N62A^ followed a negative temperature-dependence (*Q*_10fpc_^N62A^ = 0.93 ± 0.02). KaiC^E69A^ also exhibited lower *f*_pc_ in response to temperature elevation (*Q*_10fpc_^E69A^ = 0.79 ± 0.11) (Fig. 5E). While the negative temperature-dependence of the P-cycle tended to be milder than that of the ATPase activity in these site-II mutants, the over-compensatory nature seen in the ATPase activity was also carried over to some extent to the KaiABC system. The P-cycle frequency of KaiC^N62A/K172A^ was positively temperature-dependent (*Q*_10fpc_^N62A/K172A^ = 1.08 ± 0.04, Fig. 5F) as was observed for its ATPase activity (Fig. 2A and 2B).

Above results were summarized in the *Q*_10atp_–*Q*_10fpc_ plot together with the previously reported mutants (Fig. 6A). A diagonal line in the plot represents a one-to-one correspondence of the temperature dependence between the ATPase activity and the P-cycle. The plots for the site-I and site-II variants examined in this study were deviated from the diagonal line due to their milder temperature dependence of the P-cycles compared to the ATPase activities. Nevertheless, the correlation over eleven datapoints including the five mutants investigated in this study, the five CII-mutants in the previous study (Furuike *et al*., 2022c; Murayama *et al*., 2011), and KaiC^WT^, was obvious (*r* = 0.68). These observations indicate that the temperature dependence of the ATPase activity is one important factor influencing that of the P-cycle, while other factors also exert an influence.

**Figure 6.**
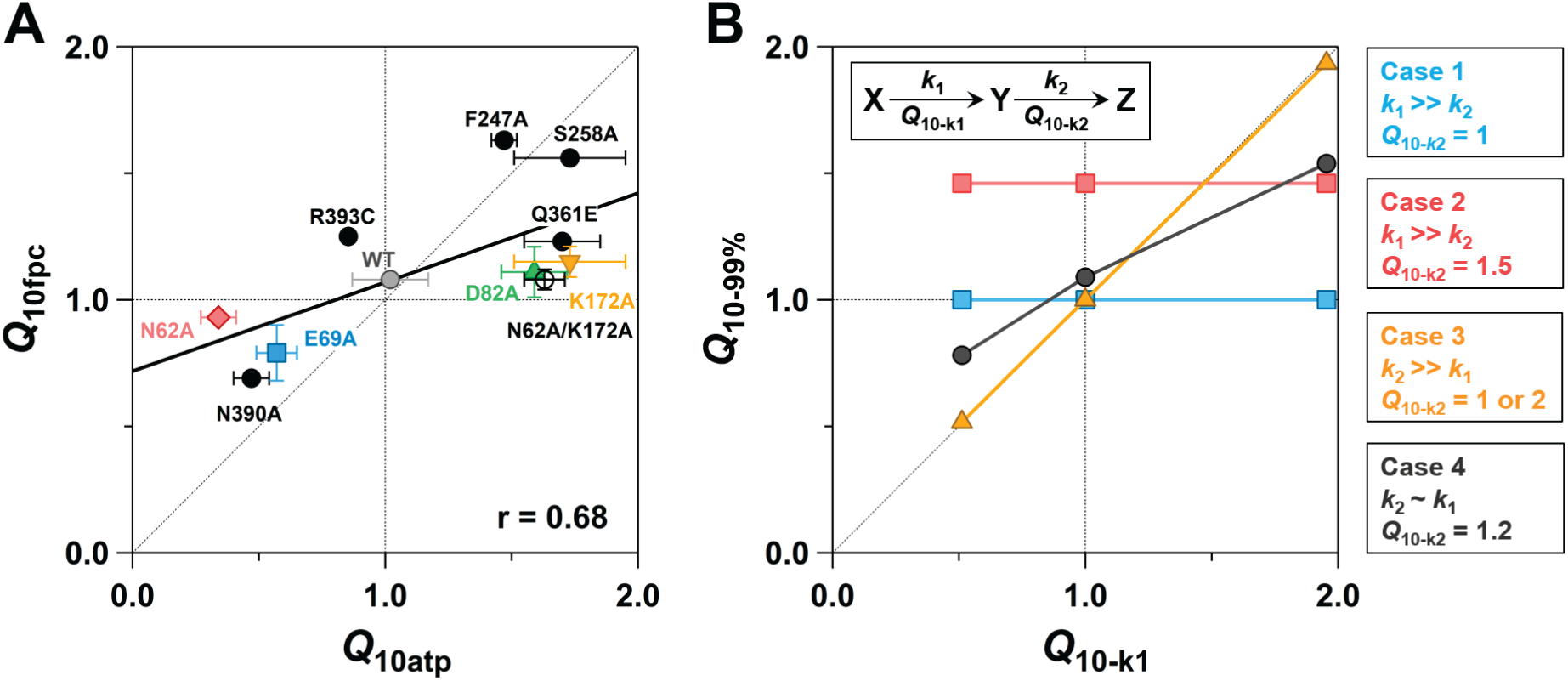
Cross-scale correlation plots of *Q*_10_. (**A**) Experimental correlation of *Q*_10_ between the P-cycle frequency (*Q*_10fpc_) and the ATPase activity (*Q*_10atp_). Error bars for *Q*_10fpc_ are small enough to be hidden behind the plots. The plot for R393C is shown without error bars (Murayama *et al*., 2011). Linear regression analysis of all the plots—including KaiC^WT^, the five mutants (colored symbols and an open circle) studied in this work, and the five mutants (black filled circles) from previous studies (Furuike *et al*., 2022c; Murayama *et al*., 2011)—yields a correlation coefficient of 0.68 (thick line). (**B**) Simulated temperature-dependence of a sequential reaction (inset). *k*_1_ and *k*_2_ represent reaction rates of the first and second steps, respectively. *τ*_99%_ denotes the time it takes for Z to reach 99% accumulation. *Q*_10-k1_, *Q*_10-k2_, and *Q*_10-99%_ indicate temperature coefficients of *k*_1_, *k*_2_, and 1/*τ*_99%_, respectively. Details of the simulation are summarized in Appendix Table S2, S3, S4, and S5.

### Local compensatory mechanism within the CI domain

Regardless of whether it is the model-A (Fig. 4) or model-B (Appendix Fig. S2), the site-I including D82 and K172 possesses the ability to locally counteract the activity increase caused by rising temperature. Thus, it is worthwhile to discuss the local role of the site-I from the perspective of its location within the CI domain. As shown in Fig. 1B, D82 and K172 are located at the CI–CI interface and are mutually stabilized by hydrogen bonds. Although the disruption of these hydrogen bonds could destabilize the ring-shaped structure, our analysis using size-exclusion chromatography with multi-angle light scattering (SEC-MALS) indicates that both ΔKaiC^D82A^ and ΔKaiC^K172A^ stably maintain the hexameric structure (Appendix Fig. S3). D82 and K172 are involved in a hydrogen-bonding network extending to R226, an Arg–finger-like residue located nearby CI-ATP (Fig. 1B). Thus, any changes in the hydrogen-bonding pattern of D82 and K172 can influence the orientation and charge of the γ-phosphate of CI-ATP via R226 (Furuike *et al*., 2022a), and also affect the positioning of a nucleophilic water molecule through a hydrogen bond with F199 (Abe *et al*., 2015) (Fig. 7A). The present site-I mutations disrupt the hydrogen bonds between D82 and K172, inevitably expanding the local conformational space of the active site within the hexamer.

**Figure 7.**
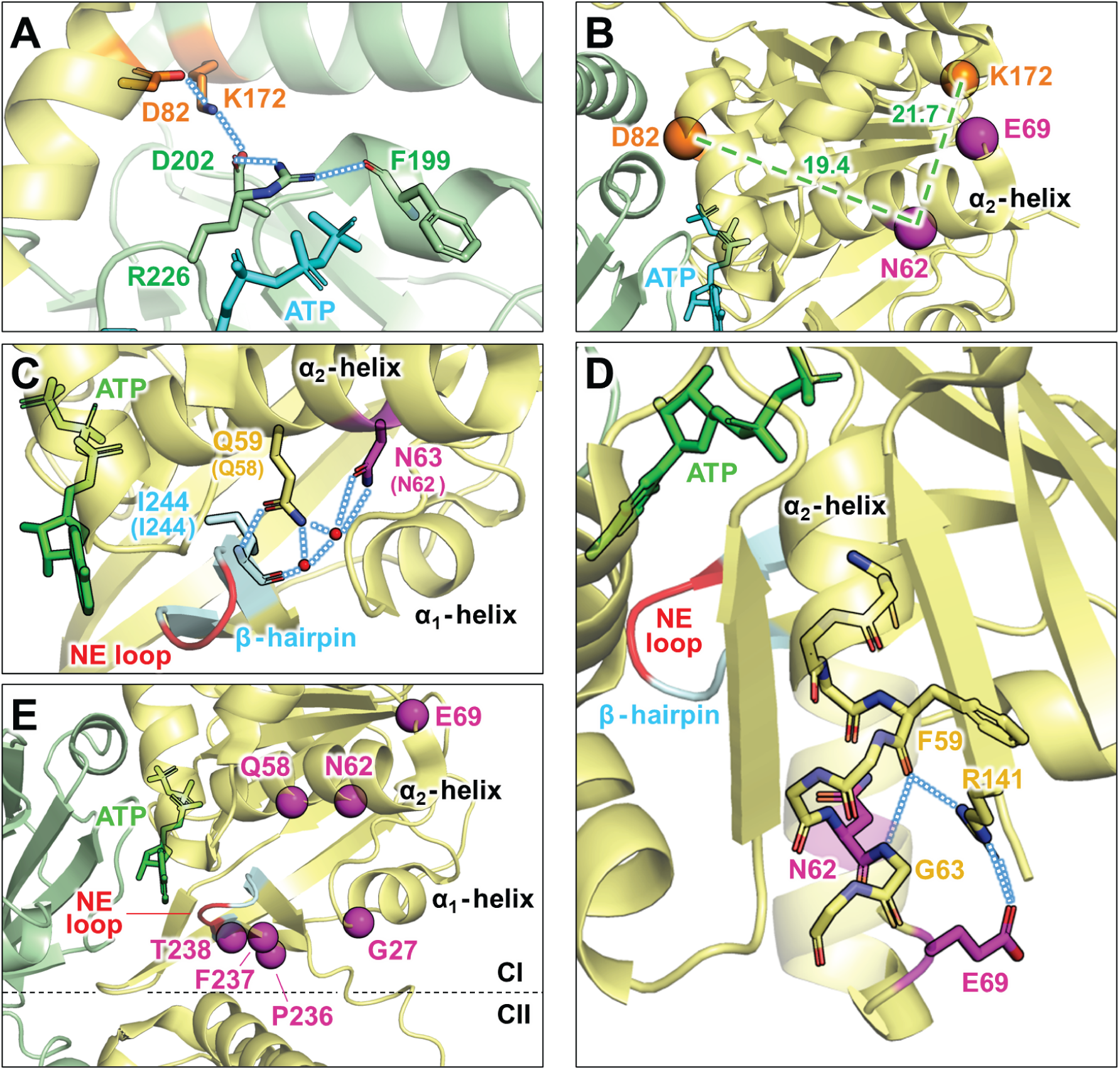
Structures of site-I and site-II affecting the temperature compensation of KaiC. (**A**) A zoomed-in-view of hydrogen-bonding network around the site-I consisting of D82 and K172. Blue dotted lines indicate potential hydrogen bonds (PDB ID: 7V3X). (**B**) Relative positions of the site-I (orange) and the site-II (magenta). The C_α_ atoms of N62, E69, D82, and K172 are presented as spheres. Green broken lines indicate distances from N62 to D82 and to K172 within the same subunit. (**C**) A zoomed-in-view of α_2_-helix and β-hairpin (cyan) in the crystal structure (PDB ID: 7DY1) of thermophilic KaiC from *Thermosynechococcus elongatus* (KaiC*^Te^*) (Furuike *et al*., 2022a). Two red spheres represent water molecules involved in a hydrogen-bonding network formed between the α_2_-helix and β-hairpin. Note that the crystal structure of KaiC*^Te^*is employed only in this panel; consequently, the residue numbers for N63 and Q59 in KaiC*^Te^* are one higher than their counterparts (in parenthesis) in KaiC^WT^ from *Synechococcus elongatus* PCC 7942. This is because, in known crystal structures of KaiC^WT^ from *Synechococcus elongatus* PCC 7942, the two water molecules are not modeled mainly due to their limited resolutions. The nucleotide enclosure (NE) loop is highlighted in red. (**D**) A hydrogen-bonding network from E69 via R141 to F59. (**E**) Mapping of C_α_ atoms (magenta spheres) of amino acid residues that exhibit negative temperature-dependences upon substitutions with alanine. *Q*_10atp_^G27A^ = 0.42, *Q*_10atp_^Q58A^ = 0.37, *Q*_10atp_^N62A^ = 0.34, *Q*_10atp_^E69A^ = 0.57, *Q*_10atp_^P236A^ = 0.62, *Q*_10atp_^F237A^ = 0.27, and *Q*_10atp_^T238A^ = 0.77 (Fig. 2B and Appendix Fig. S4).

We emphasize that a more pronounced increase in the local conformational space at lower temperatures leads to an increase in *Q*_10_. Here, we assume that the number of possible local conformations of the active site is *N*, and that one of *N* corresponds to a reaction-ready state. In this case at low temperatures, the probability of selecting the active conformation is 1/*N*. Using a frequency (*ω*) of that selection, the reaction rate per unit time becomes proportional to *ω*/*N*. At high temperatures, increases in both the number of the conformations (*N*+δ*N*) and selection frequency (*ω*+δ*ω*) occur; therefore, the reaction rate is expressed as (*ω*+δ*ω*)/(*N*+δ*N*). A ratio of the reaction rate at high temperatures to that at low temperatures is,

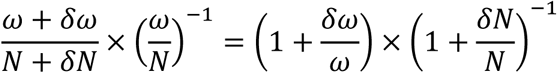

where the increase (or decrease) in the selection frequency is counterbalanced by the increase (or decrease) in the number of the possible local conformations. Since the disruption of the hydrogen bonds is one primary factor enhancing structural flexibility, the number of the possible conformations increases in the site-I mutants; e.g., additional *n* conformations at both low and high temperatures. In this case, the ratio of the reaction rates is increased:

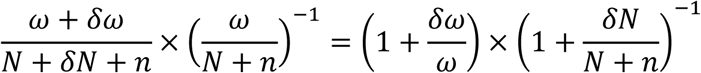

We suggest that the hydrogen bonds in the site-I contribute to the local compensatory regulation by narrowing the local conformational space of the active site (*n* ∼ 0).

### Non-local compensatory mechanism through CI-CII domain coordination

One of the differences between the model-A and model-B lies in whether the inhibition by the site-II, which prevents the ATPase activity from becoming over-compensated, acts on the non-local compensatory pathway mediated by the site-I (model-B in Appendix Fig. S2) or on the separate route that does not pass through the site-I (model-A in Fig. 4). One reason why the model-B appears less probable is that the site-II is not close enough to the site-I to exert the inhibitory effect on it. The site-II is positioned approximately 20 Å away from the site-I (Fig. 7B), and the structural causality between the two sites is not evident from the known structures of KaiC.

On the contrary, the model-A predicts that the non-local compensatory regulation passes through a vicinity of the site-II. One such candidate is the α_2_-helix in the CI domain. CI-ATP interacts with a phosphate-binding loop (P-loop) located at the N-terminus of the α_2_-helix, whereas both N62 and E69 are placed on its C-terminal side (Fig. 1C). The side chain of N62 interacts with a β-hairpin harboring a nucleotide enclosure (NE) loop (Mukaiyama *et al*, 2025) through a hydrogen-bonding network mediated by two water molecules and Q58 (Fig. 7C). Considering the proximity of this β-hairpin to the CII domain, we propose that the non-local compensatory regulation lies on a pathway from the CII domain to the NE loop via the β-hairpin. In this case, a scenario emerges in which the α_2_-helix interacts with the β-hairpin to prevent the NE loop from being excessively affected when the CII domain approaches (Chang *et al*, 2012; Swan *et al*, 2022).

E69 can also indirectly influence the interaction between α_2_-helix and the β-hairpin. A hydrogen-bonding network originating from the side chain of E69 extends via a side chain of R141 to a carbonyl oxygen of the main chain of F59 forming the α-helical structure (Fig. 7D). The hydrogen bond between the side chain of R141 and the main chain of F59 is unusual, as carbonyl oxygens on the α-helical main chain regularly form hydrogen bonds exclusively with amide hydrogens located four residues downstream. The unusual hydrogen-bonding pattern of F59 is partially attributed to the fact that a residue at the 63rd position is Gly (Fig. 7D). As judged by the hydrogen-bonding distance, the carbonyl oxygen of F59 receives stronger proton donation from an N_η_H_η_ moiety of R141 (2.7 Å) than from the amide proton of G63 (3.3 Å). Therefore, the E69A substitution destabilizes the α_2_-helix or at least its C-terminal part containing N62 by increasing the positive charge at R141 and indirectly strengthening the hydrogen bond between F59 and R141.

If our interpretation on the site-II and the non-local compensatory pathway is plausible, we would expect to find mutations exhibiting the over-compensated ATPase activity widely around the α_2_-helix and the β-hairpin. In fact, the negative temperature-dependences of the ATPase activity were observed in other surrounding regions (Fig. 7E and Appendix Fig. S4); the α_1_-helix (KaiC^G27A^), the β-hairpin (KaiC^P236A^, KaiC^F237A^, and KaiC^T238A^), and the interaction site between the α_2_-helix and β-hairpin (KaiC^Q58A^). This means that the site-II is not necessarily limited to the specific residues such as N62 and E69. Further experimental approaches are needed to capture transient structural changes around the site-II and to quantify the temperature dependence of their frequency and magnitude.

### Similarities to temperature-compensation mechanisms in other organisms

The ATPase activity of KaiC has been considered as one of the key factors affecting the properties of the KaiABC system. Recently, Sasai *et al*. developed a theoretical model of the KaiABC system, which reproduces temperature-compensated circadian oscillations at an ensemble level (Sasai & Fujishiro, 2026). They treated each KaiC hexamer as an oscillator in which the CI-ATPase cycle and the phosphorylation state of the CII domain are coupled via the CI-CII interface, and treated multiple hexamers as being coupled through their inter-molecular interactions. Interestingly, by taking into account an entropy production resulting from the dissipation of free energy supplied by the ATP hydrolysis in each cycle, they derived a relationship between the ATP consumption by KaiC and *f*_pc_. The equation (17) in their paper predicts that the correlation line on the *Q*_10atp_–*Q*_10fpc_ plot rotates clockwise with a slope of approximately 0.5. Their result shows a reasonable agreement with our experimental slope of 0.35 ±0.13 (Fig. 6A).

The correlation coefficient of less than unity in Fig. 6A can be also explained by influences of other factors. We investigated the temperature dependence of a three-component sequential reaction (X → Y → Z) at both the system-wide and individual-step levels (Fig. 6B, and Appendix Table S2, S3, S4, and S5). We defined *τ*_99%_ as the time it takes for the component Z to reach 99% accumulation and calculated the temperature coefficient of 1/*τ*_99%_ (*Q*_10-99%_). Under conditions where a rate of the first step (*k*_1_: X → Y) was much larger than a rate of the second step (*k*_2_: Y → Z), *Q*_10-99%_ was always equal to a temperature coefficient of the second step (*Q*_10-k2_) regardless of that of the first step (*Q*_10-k1_) (case 1 and 2 in Fig. 6B). Diagonal correlations emerged between *Q*_10-99%_ and *Q*_10-k1_ irrespective of *Q*_10-k2_, when *k*_2_ significantly exceeded *k*_1_ (case 3 in Fig. 6B). It was possible to reproduce a clockwise rotation of the correlation line under a specific condition, where *k*_1_ was comparable to *k*_2_ with *Q*_10-k2_ of approximately 1.2 (case 4 in Fig. 6B). These observations indicate that, in the KaiABC system, the ATPase cycle of KaiC is kinetically coupled to another process that proceeds at a comparable rate in a reasonably temperature-compensated manner.

It is important to discuss our observations with respect to temperature-compensation mechanisms in other organisms. In mammalian clock systems, the temperature compensation has been studied from the perspective of the phosphorylation of clock proteins by casein kinase I delta (CKIδ). An earlier study reported that the CKIδ-dependent phosphorylation of model substrates proceeds on the order of minutes in a temperature-compensated manner (Isojima *et al*, 2009). A subsequent study demonstrated that the temperature compensation of the CKIδ activity results from a reversal in the temperature dependence of its affinities against for substrates and products (Shinohara *et al*., 2017). While the introduction of K224D substitution into CKIδ causes its *Q*_10_ to rise to approximately 2.0, the *Q*_10_ value of the clock frequency remains approximately 1.1. The study shares several similarities with our observations, particularly in that the temperature sensitivity or insensitivity at the enzymatic level can influence that at the system-wide level, and that this effect is sometimes observed to be attenuated depending on the nature of the system.

In *Arabidopsis* circadian clock, two proteins known as TOC1 and PRR5 possess a compensatory function that counteracts the frequency increase at higher temperatures (Maeda *et al*., 2024). Interestingly, to prevent the compensatory effects of TOC1 and PPR5 from becoming excessive, their accumulation is downregulated by interactions with a component of the ubiquitin ligase, LKP2. Although the difference lies in whether it is intra-molecular or inter-molecular scale, the circadian clocks of cyanobacteria and *Arabidopsis* share a common feature: the coexistence of one factor suppressing another factor that counteracts the acceleration caused by rising temperature. This may be the basic structure of the temperature-compensation network that is conserved across scales and species.

## Methods

### Expression and purification

KaiC^WT^ and all ΔKaiCs were expressed as glutathione S-transferase (GST)-tagged forms using a pGEX-6P-1 plasmid (Novagen) and *Escherichia coli* (*E. coli*) BL21(DE3) (Nishiwaki *et al*., 2007). Other KaiC mutants were expressed as C-terminal histidine (His)-tagged forms using a pET-3a plasmid (Novagen) and BL21(DE3)pLysE (Ouyang *et al*, 2019). The GST-tagged forms were overexpressed at 37°C in the presence of 0.04 mg/mL ampicillin through a 6-hour pre-culture (Lysogeny Broth (LB), 200 mL) followed by an 18-hour main culture (Terrific Broth (TB), 800 mL × 4). In the case of the His-tagged forms, cells were grown at 37°C though a 16-hour pre-culture (LB, 10 mL) in the presence of 0.04 mg/mL ampicillin and 0.035 mg/mL chloramphenicol and a subsequent main culture (TB, 400 mL × 1) in the presence of 0.04 mg/mL ampicillin. The His-tagged proteins were overexpressed at 37°C through a 5-hour main culture after the addition of isopropyl-β-D-thiogalactopyranoside at a final concentration of 0.05 mM. The cells were collected by centrifuge and stored at -25°C.

The cell pellets were suspended into a buffer containing 20 mM Tris-HCl (pH 8.0), 150 mM NaCl, 5 mM MgCl_2_, 1 mM DTT, 0.5 mM EDTA, 1 mM ATP, and 40 U/mL Benzonase nuclease (Merck) and crushed by sonication. After stirring for 1 h at 4°C in the presence of 1% TritonX-100, the lysate was centrifuged to remove insoluble fraction. In the case of the GST-tagged forms, the supernatant was applied to a glutathione sepharose column (Cytiva) and the GST-tagged protein was selectively eluted after cleaving its GST-tag with 53 U/mL PreScission protease (Cytiva). In the case of the His-tagged forms, the supernatant was applied to a Ni-sepharose column and the His-tagged protein was eluted with a buffer containing 480 mM imidazole, 20 mM Tris-HCl (pH 7.4), 500 mM NaCl, 1 mM ATP, and 5 mM MgCl_2_. In each case, the affinity-purified sample was further purified using a size-exclusion column (Sephacryl S300 for the GST-tagged forms and Superdex 200 increase 10/300 GL for the His-tagged forms) and an anion-exchange column (Resource Q, Cytiva) as described previously (Mukaiyama *et al*, 2019; Ouyang *et al*., 2019). The purified samples were finally dissolved in a Tris-buffer (20 mM Tris-HCl, 150 mM NaCl, 5 mM MgCl_2_, 1 mM DTT, 0.5 mM EDTA, 1 mM ATP, and pH 8.0) and stored at -80°C.

### ATPase activity measurements

KaiC was dissolved in the Tris-buffer at concentrations of 0.2–0.5 mg/mL and incubated at different temperatures (25, 30, and 35°C). Accumulated ADP was separated from ATP on reversed-phase column (InertSustain C18, GL Sciences) at a flow rate of 0.4 mL/min with a mobile phase of 16% (v/v) acetonitrile, 20 mM ammonium phosphate, and 10 mM tetrabutylammonium hydrogen sulfate (pH 8.5) (Murayama *et al*., 2011; Ouyang *et al*., 2019) using high performance liquid chromatography system (EXTREMA, JASCO). At each time point, the concentration of ADP was determined from its peak area (Murayama *et al*., 2011; Ouyang *et al*., 2019). ATPase activity was defined as the number of ADP produced by one KaiC molecule in a monomer unit per day. Since it takes full-length KaiCs approximately 5 h to reach their steady states after the samples were transferred from on ice to incubation temperatures, the data points earlier than the incubation time of 5 h were excluded from the analysis.

### *In vitro* phosphorylation cycle assay

KaiA, KaiB, and KaiC were mixed in the Tris-buffer at concentrations of 0.04 mg/mL, 0.04 mg/mL, and 0.2 mg/mL, respectively. The mixture was incubated at different temperatures (30, 35, and 40°C) and aliquots were taken at several-hour interval. Phosphorylated and non-phosphorylated KaiCs were separated by sodium dodecyl sulfate polyacrylamide gel electrophoresis (Nakajima *et al*., 2005) and the ratio of phosphorylated KaiC at each time point was quantified by densitometric analysis of gel bands as described previously (Furuike *et al*., 2016). To estimate the period length, the time evolution of the fraction of the phosphorylated KaiC was fitted to a cosine function (Furuike *et al*., 2016).

### Estimation of *Q*_10_ values

Temperature dependences of the ATPase activity and the P-cycle frequency were subjected to Arrhenius plot analysis. The resultant *E*_A_ was converted to the *Q*_10_ value at 30°C using the following equation (Furuike *et al*., 2016; Furuike *et al*., 2022c),

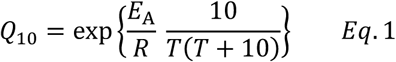

where *T* and *R* represent absolute temperature and the gas constant, respectively

### Size-exclusion chromatography with multi-angle light scattering (SEC-MALS)

ΔKaiC^WT^, ΔKaiC^D82A^, and ΔKaiC^K172A^ were dissolved into the Tris-buffer at concentrations of 0.22 mg/mL, 0.27 mg/mL, and 0.25 mg/mL, respectively. 500 µL of samples was subjected to the size-exclusion column (Superdex 200 increase 10/300 GL, Cytiva) combined with a multi-angle static light scattering system (Viscoteck TDA305, Malvern). The samples were eluted at a flow rate of 0.4 mL/min at 4°C and the scattered light intensity at angles of 7° and 90° were simultaneously monitored to estimate molecular masses at the eluted peaks.

### X-ray crystallography

Crystals of ΔKaiC^N62A^ and ΔKaiC^E69A^ were obtained by the sitting-drop vapor diffusion method. ΔKaiC^N62A^ and ΔKaiC^E69A^ at 5 mg/mL were mixed with the reservoir solution containing 100 mM Tris-HCl (pH 8.5), 130-160 mM MgSO_4_, 31-34 % (w/v) PEG400, and 5 mM adenylyl imidodiphosphate (AMP-PNP), and incubated at 30°C. The crystals were frozen in liquid nitrogen.

X-ray diffraction data were collected at 100 K under a cryo-stream on beamline BL44XU at SPring-8 (Harima, Japan). Diffraction images were recorded using EIGER X 16M detector (Dectris) and screened with the semi-automatic data-processing pipeline, KAMO (Yamashita *et al*, 2018). Diffraction intensity data were processed using XDS (Kabsch, 2010). The initial phases for both ΔKaiC^N62A^ and ΔKaiC^E69A^ were obtained by molecular replacement using Phaser-MR in PHENIX program package (version 2.1-6048) (Adams *et al*, 2010) and ΔKaiC^WT^ structure (PDB ID: 4TL7) (Abe *et al*., 2015). Refinement and model building were conducted using PHENIX with R-free flags transferred from 4TL7 and Coot (version 0.9.8.95) (Emsley *et al*, 2010), respectively. Graphic representations were generated using PyMOL (Schrödinger) (version3.0.2). The statistics for data collection and refinement are listed in Appendix Table S1.

## Data availability

The atomic coordinates of ΔKaiC^N62A^ (PDB ID: 26QX) and ΔKaiC^E69A^ (PDB ID: 26RC) have been deposited in the Protein Data Bank.

## Author contributions

**Kanta Kondo**: Data curation; Formal analysis; Validation; Investigation, Visualization; Writing—original draft; Writing—review and editing. **Yoshihiko Furuike**: Data curation; Formal analysis; Validation; Visualization; Project administration, Funding acquisition; Writing—review and editing. **Kota Horiuchi**: Formal analysis; Validation; Methodology. **Yasuhiro Onoue**: Formal analysis; Validation; Methodology. **Eiki Yamashita**: Validation; Methodology. **Shuji Akiyama**: Conceptualization; Project administration, Funding acquisition; Validation; Visualization; Writing—original draft; Writing—review and editing.

## Disclosure and competing interests statement

The authors declare that they have no conflict of interest.

## Acknowledgements

This study was supported by JSPS Grants-in-Aid for Scientific Research (22H04984 and 24H02301 to S.A., and 26H01830 to Y.F.) and partly by Takeda Science Foundation (to S.A.) and Toyoaki Scholarship Foundation (to S.A.). X-ray diffraction data were collected at beamline BL44XU at the SPring-8 facility under the proposals 2023A6500, 2024A6930, 2024B6930, and 2025A6526.

## Notes

### Competing Interest Statement

The authors have declared no competing interest.

