## Appendix for "Local and non-local impacts of intra-hexamer interactions on temperature compensation of KaiC"

**Contents:**

Appendix Figure S1 Page 2
Appendix Figure S2 Page 3
Appendix Figure S3 Page 4
Appendix Figure S4 Page 5
Appendix Table S1 Page 6
Appendix Table S2 Page 7
Appendix Table S3 Page 8
Appendix Table S4 Page 9
Appendix Table S5 Page 10

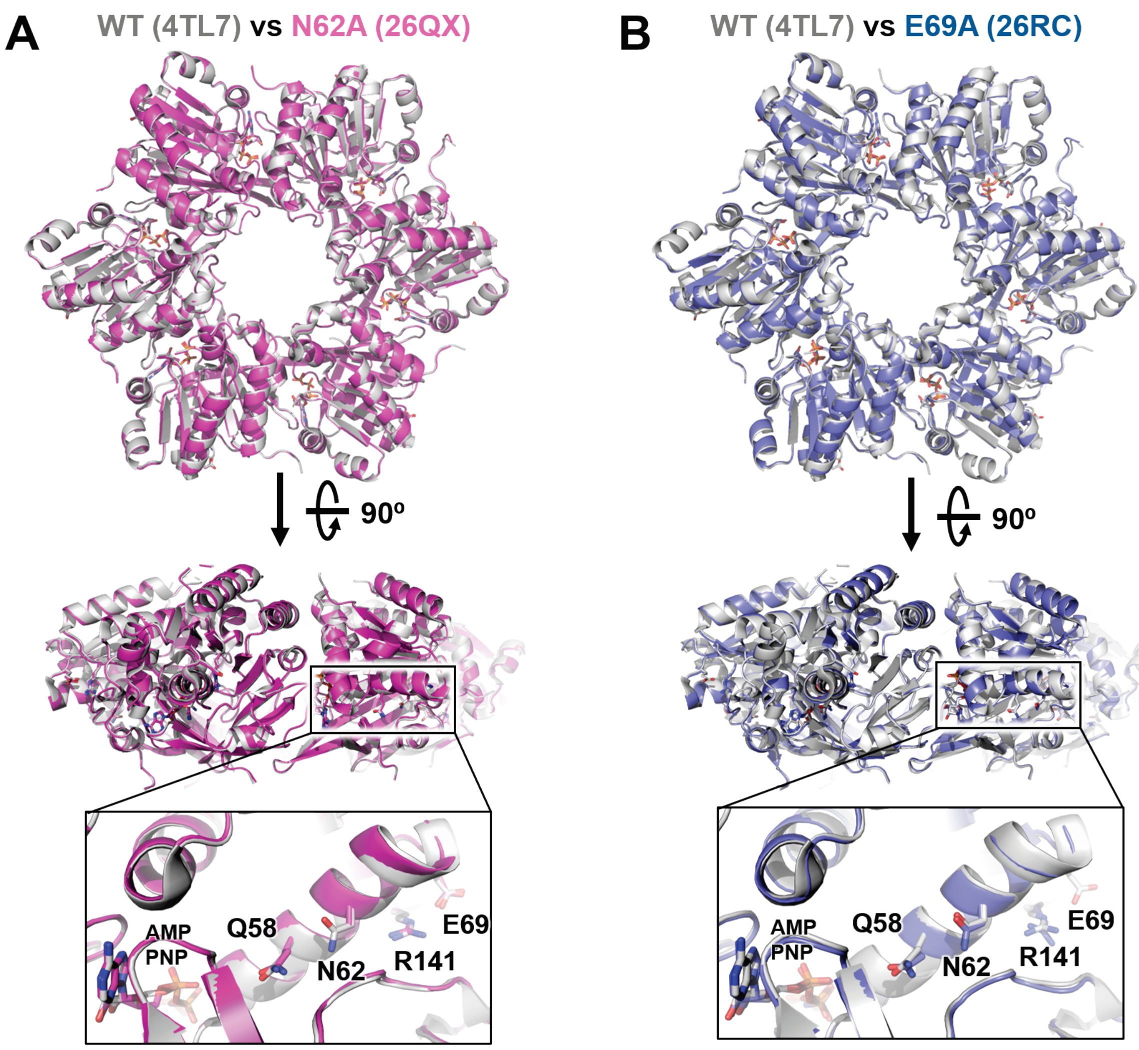


**Appendix Figure S1. Crystal structures of ΔKaiCs.** (**A**) Superimposition of $\Delta KaiC^{WT}$ (gray, PDB ID: 4TL7, 1.9 Å resolution) and $\Delta KaiC^{N62A}$ (magenta, PDB ID: 26QX, 2.2 Å resolution). The main-chain root-mean-squared deviation (RMSD) values were 0.37 Å and 0.24 Å for hexameric and monomeric superimpositions, respectively. (**B**) Superimposition of $\Delta KaiC^{WT}$ (gray, PDB ID: 4TL7) and $\Delta KaiC^{E69A}$ (blue, PDB ID: 26RC, 2.8 Å resolution). The main-chain RMSD values were 0.56 Å and 0.39 Å for hexameric and monomeric superimpositions, respectively.

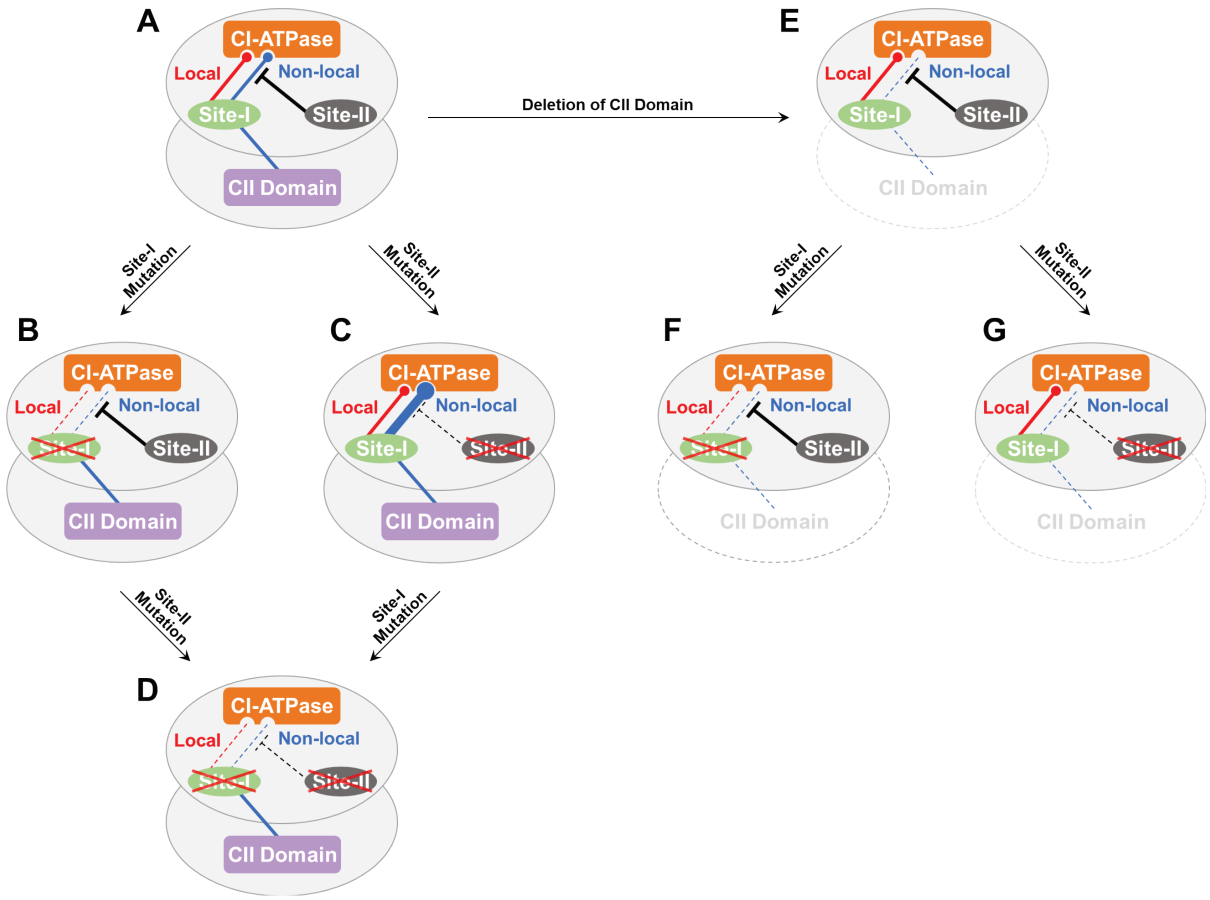


**Appendix Figure S2. A conceptual illustration of model-B.** A site-I acts on the CI-ATPase cycle through a local temperature-compensatory regulation (red-tipped arrows). The site-I receives a non-local compensatory regulation from the CII domain and forwards it to the CI-ATPase cycle (blue-tipped arrows). A site-II inhibits only the non-local compensatory pathway from the site-I to the CI-ATPase cycle (black inhibitory arrows). (**A**) KaiC$^{WT}$. (**B**) Site-I mutants of KaiC$^{WT}$. Both the local and non-local compensatory regulations are absent or considerably weakened. (**C**) Site-II mutants of KaiC$^{WT}$. The non-local compensatory regulation is excessively enhanced (thick blue-tipped arrow) by the release of inhibition by the site-II, resulting in the over-compensated CI-ATPase. (**D**) Site-I/II double mutant of KaiC$^{WT}$. (**E**) ΔKaiC$^{WT}$. The deletion of the CII domain results in a loss or significant attenuation of the non-local compensatory regulation. (**F**) Site-I mutants of ΔKaiC$^{WT}$. (**G**) Site-II mutants of ΔKaiC$^{WT}$. The CI-ATPase activity is yet under some local compensatory regulation.

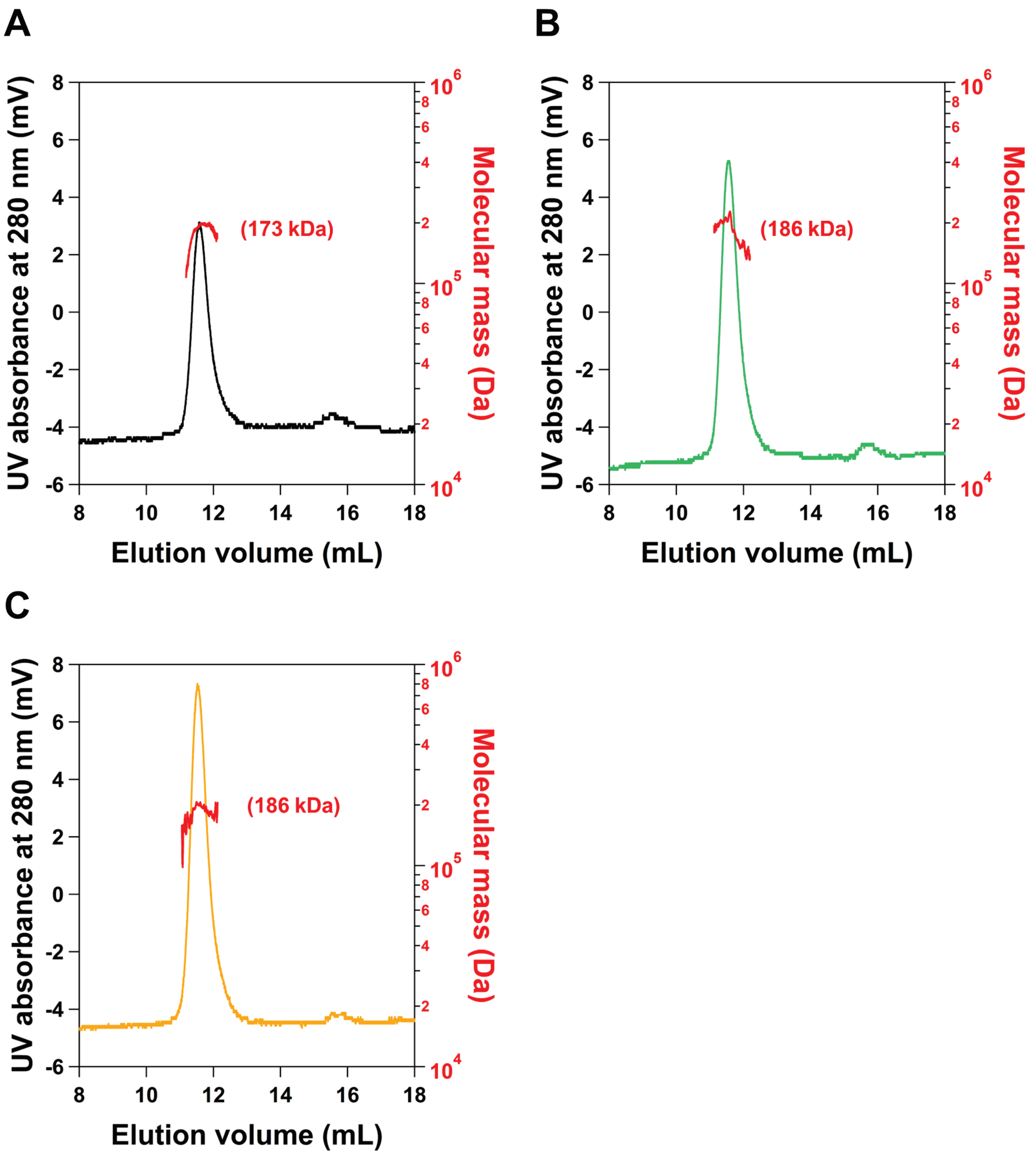


**Appendix Figure S3. SEC-MALS analysis of site-I mutants.** SEC chromatograms and estimated molecular masses of (**A**) ΔKaiC$^{WT}$, (**B**) ΔKaiC$^{D82A}$, and (**C**) ΔKaiC$^{K172A}$. The theoretical molecular mass of ΔKaiC$^{WT}$ hexamer is 171,762 Da. Values in parentheses represent estimated weight-averaged molecular masses at the peak.

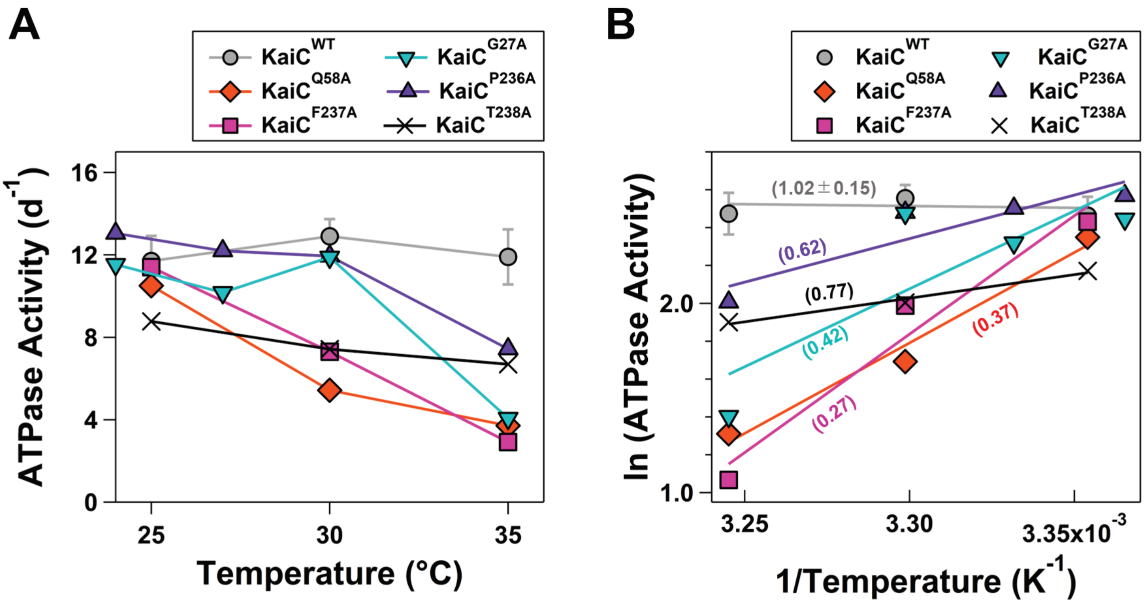


**Appendix Figure S4. Temperature dependence of the ATPase activity of site-II related mutants.** (**A**) ATPase activities and (**B**) Arrhenius-plot analysis of the site-II related mutants. Values in parentheses represent $Q_{10atp}$. The samples sizes (n): n = 1 except for KaiC$^{WT}$ (n = 3).

**Appendix Table S1. Data collection and refinement statistics.** [a]Values in parentheses correspond to the highest-resolution shell. [b]$R_{merge} = \Sigma|I - \langle I \rangle| / \Sigma I$, where $I$ corresponds to the observed intensity of reflections. [c]$R_{work,\ free} = \Sigma(|F_{obs}| - |F_{calc}|) / \Sigma|F_{obs}|$. $R_{free}$ is the cross-validation of the $R$-factor using the test reflections, 5% of the data, not included in the refinements.

| **Protein** | **$\Delta$KaiC$^{N62A}$** | **$\Delta$KaiC$^{E69A}$** |
|---|---|---|
| **X-ray Source** | SPring-8 BL44XU | SPring-8 BL44XU |
| **Data Collection** | | |
| Space group | $P3_121$ | $P3_121$ |
| Unit cell length *a, b, c* (Å) | 107.6, 107.4, 225.9 | 107.4, 107.4, 218.8 |
| Unit cell angle α, β, γ (°) | 90, 90, 120 | 90, 90, 120 |
| Wavelength (Å) | 0.9 | 0.9 |
| Resolution range (Å)[a] | 48.59-2.18 (2.26-2.18) | 48.22-2.79 (2.89-2.79) |
| Total reflections | 807631 (74898) | 371506 (36231) |
| Unique reflections | 79889 (7779) | 37112 (3578) |
| Redundancy | 10.11 (9.63) | 10.01 (10.13) |
| Completeness (%) | 99.86 (98.76) | 99.55 (97.39) |
| (I)/sigma(I) | 16.4 (2.6) | 13.0 (2.4) |
| $R_{merge}$[b] | 0.079 (0.514) | 0.140 (0.908) |
| **Model building** | | |
| Molecular replacement | 4TL7 | 4TL7 |
| Total atoms | 11067 | 9811 |
| Protien | 10431 | 9604 |
| Ligands | 218 | 196 |
| Water | 418 | 11 |
| $R_{work}$ (%)[c] | 17.9 | 23.8 |
| $R_{free}$ (%)[c] | 21.8 | 29.0 |
| R.M.S.D. from ideality | | |
| Bond length (Å) | 0.004 | 0.004 |
| Bond angles (°) | 0.68 | 0.77 |
| Average B factors (Å$^2$) | 48.6 | 82.1 |
| Ramachandran plot | | |
| Favored (%) | 97.1 | 97.2 |
| Allowed (%) | 2.9 | 2.8 |
| Outliers (%) | 0.0 | 0.0 |
| **PDB code** | **26QX** | **26RC** |

**Appendix Table S2. Parameters used for simulating Case 1 of the sequential reaction shown in Fig. 6B.**

<table>
<tr><th>$T$ (°C)</th><th>$1/(T+273.15)$ ($K^{-1}$)</th><th>$k_1$</th><th>$Q_{10\text{-k1}}$</th><th>$k_2$</th><th>$Q_{10\text{-k2}}$</th><th>$1/\tau_{99\%}$</th><th>$Q_{10\text{-}99\%}$</th></tr>
<tr><td>20</td><td>0.00341</td><td>2000</td><td rowspan="3">0.51</td><td rowspan="9">1.00</td><td rowspan="9">1.00</td><td>0.2083</td><td rowspan="3">1.00</td></tr>
<tr><td>30</td><td>0.00330</td><td>1000</td><td>0.2083</td></tr>
<tr><td>40</td><td>0.00319</td><td>500</td><td>0.2083</td></tr>
<tr><td>20</td><td>0.00341</td><td>1000</td><td rowspan="3">1.00</td><td>0.2083</td><td rowspan="3">1.00</td></tr>
<tr><td>30</td><td>0.00330</td><td>1000</td><td>0.2083</td></tr>
<tr><td>40</td><td>0.00319</td><td>1000</td><td>0.2083</td></tr>
<tr><td>20</td><td>0.00341</td><td>500</td><td rowspan="3">1.95</td><td>0.2083</td><td rowspan="3">1.00</td></tr>
<tr><td>30</td><td>0.00330</td><td>1000</td><td>0.2083</td></tr>
<tr><td>40</td><td>0.00319</td><td>2000</td><td>0.2083</td></tr>
</table>

**Appendix Table S3. Parameters used for simulating Case 2 of the sequential reaction shown in Fig. 6B.**

| $T$ (°C) | $1/(T+273.15)$ ($K^{-1}$) | $k_1$ | $Q_{10\text{-k1}}$ | $k_2$ | $Q_{10\text{-k2}}$ | $1/\tau_{99\%}$ | $Q_{10\text{-}99\%}$ |
|---|---|---|---|---|---|---|---|
| 20 | 0.00341 | 2000 | | 0.67 | | 0.1429 | |
| 30 | 0.00330 | 1000 | 0.51 | 1.00 | | 0.2083 | 1.46 |
| 40 | 0.00319 | 500 | | 1.50 | | 0.3125 | |
| 20 | 0.00341 | 1000 | | 0.67 | | 0.1429 | |
| 30 | 0.00330 | 1000 | 1.00 | 1.00 | 1.48 | 0.2083 | 1.46 |
| 40 | 0.00319 | 1000 | | 1.50 | | 0.3125 | |
| 20 | 0.00341 | 500 | | 0.67 | | 0.1429 | |
| 30 | 0.00330 | 1000 | 1.95 | 1.00 | | 0.2083 | 1.46 |
| 40 | 0.00319 | 2000 | | 1.50 | | 0.3125 | |

**Appendix Table S4. Parameters used for simulating Case 3 of the sequential reaction shown in Fig. 6B.**

| $T$ (°C) | $1/(T+273.15)$ ($K^{-1}$) | $k_1$ | $Q_{10\text{-k1}}$ | $k_2$ | $Q_{10\text{-k2}}$ | $1/\tau_{99\%}$ | $Q_{10\text{-}99\%}$ |
|---|---|---|---|---|---|---|---|
| 20 | 0.00341 | 2.00 | | 1000 | | 0.4167 | |
| 30 | 0.00330 | 1.00 | 0.51 | 1000 | | 0.2083 | 0.52 |
| 40 | 0.00319 | 0.50 | | 1000 | | 0.1064 | |
| 20 | 0.00341 | 1.00 | | 1000 | | 0.2083 | |
| 30 | 0.00330 | 1.00 | 1.00 | 1000 | 1.00 | 0.2083 | 1.00 |
| 40 | 0.00319 | 1.00 | | 1000 | | 0.2083 | |
| 20 | 0.00341 | 0.50 | | 1000 | | 0.1064 | |
| 30 | 0.00330 | 1.00 | 1.95 | 1000 | | 0.2083 | 1.93 |
| 40 | 0.00319 | 2.00 | | 1000 | | 0.4167 | |
| 20 | 0.00341 | 2.00 | | 500 | | 0.4167 | |
| 30 | 0.00330 | 1.00 | 0.51 | 1000 | | 0.2083 | 0.52 |
| 40 | 0.00319 | 0.50 | | 2000 | | 0.1064 | |
| 20 | 0.00341 | 1.00 | | 500 | | 0.2083 | |
| 30 | 0.00330 | 1.00 | 1.00 | 1000 | 1.95 | 0.2083 | 1.00 |
| 40 | 0.00319 | 1.00 | | 2000 | | 0.2083 | |
| 20 | 0.00341 | 0.50 | | 500 | | 0.1064 | |
| 30 | 0.00330 | 1.00 | 1.95 | 1000 | | 0.2083 | 1.93 |
| 40 | 0.00319 | 2.00 | | 2000 | | 0.4167 | |

**Appendix Table S5. Parameters used for simulating Case 4 of the sequential reaction shown in Fig. 6B.**

| $T$ (°C) | $1/(T+273.15)$ ($K^{-1}$) | $k_1$ | $Q_{10\text{-k1}}$ | $k_2$ | $Q_{10\text{-k2}}$ | $1/\tau_{99\%}$ | $Q_{10\text{-99\%}}$ |
|---|---|---|---|---|---|---|---|
| 20 | 0.00341 | 2.00 | | 0.83 | | 0.1613 | |
| 30 | 0.00330 | 1.00 | 0.51 | 1.00 | | 0.1471 | 0.78 |
| 40 | 0.00319 | 0.50 | | 1.20 | | 0.0962 | |
| 20 | 0.00341 | 1.00 | | 0.83 | | 0.1351 | |
| 30 | 0.00330 | 1.00 | 1.00 | 1.00 | 1.20 | 0.1471 | 1.09 |
| 40 | 0.00319 | 1.00 | | 1.20 | | 0.1613 | |
| 20 | 0.00341 | 0.50 | | 0.83 | | 0.0893 | |
| 30 | 0.00330 | 1.00 | 1.95 | 1.00 | | 0.1471 | 1.54 |
| 40 | 0.00319 | 2.00 | | 1.20 | | 0.2174 | |
